# Cryopreservation of Paramecium bursaria Chlorella Virus-1 during an active infection cycle of its host

**DOI:** 10.1101/528786

**Authors:** Samantha R. Coy, Alyssa N. Alsante, James L. Van Etten, Steven W. Wilhelm

**Affiliations:** Department of Microbiology, University of Tennessee, Knoxville, Tennessee, United States of America; Department of Plant Pathology and Nebraska Center for Virology, University of Nebraska, Lincoln, Nebraska, United States of America

## Abstract

Best practices in laboratory culture management often include cryopreservation of microbiota, but this can be challenging with some virus particles. By preserving viral isolates researchers can mitigate genetic drift and laboratory-induced selection, thereby maintaining genetically consistent strains between experiments. To this end, we developed a method to cryopreserve the model, green-alga infecting virus, *Paramecium bursaria Chlorella virus 1* (PBCV-1). We explored cryotolerance of the infectivity of this virus particle, whereby freezing without cryoprotectants was found to maintain the highest infectivity (~2.5%). We then assessed the cryopreservation potential of PBCV-1 during an active infection cycle in its *Chlorella variabilis* NC64A host, and found that virus survivorship was highest (69.5 ± 16.5 %) when the infected host is cryopreserved during mid-late stages of infection (*i.e*., coinciding with virion assembly). The most optimal condition for cryopreservation was observed at 240 minutes post-infection. Overall, utilizing the cell as a vehicle for viral cryopreservation resulted in 24.9 – 30.1 fold increases in PBCV-1 survival based on 95% confidence intervals of frozen virus particles and virus cryopreserved at 240 minutes post-infection. Given that cryoprotectants are often naturally produced by psychrophilic organisms, we suspect that cryopreservation of infected hosts may be a reliable mechanism for virus persistence in non-growth permitting circumstances in the environment, such as ancient permafrosts.

## Introduction

Viruses are abundant components of all biological systems and they likely infect every lineage of eukaryotic algae. Their impact is most readily noticed following infection and lysis of abundant bloom forming algae (1–3), though lytic activity of all algal viruses contributes to significant biomass recycling *via* the ‘viral shunt’ (4). To date, 65 eukaryotic algal viruses have been isolated and developed as laboratory strains (5, 6). Most of these are maintained through serial propagation on their respective hosts. Though this has been effective for culturing many strains over the last few decades (7, 8), each passage allows for genetic mutations that can accumulate in a population (9), leading to a deviation from a standard ‘wild-type.’ Moreover, it is imperative to control evolution following the development of genetically tractable algal hosts (10) and (ultimately) virus systems. A protocol for successful virus cryobiological preservation would offer an opportunity to maintain genotypically consistent virus stocks.

Cryopreservation is not a new concept in biological sciences. For most protocols, it involves controlled cooling of biota to sub-freezing temperatures to achieve biological cessation while preserving viability. This most often manifests as slow-cooling at a rate of 1° C / min in the presence of osmoprotectant(s) (*e.g*., dimethylsulfoxide (DMSO), glycerol) for long-term storage at −130° C or below (11). Too slow a cooling rate can result in higher intracellular concentration of osmoprotectants, resulting in toxicity, whereas too fast a cooling rate allows the formation of intracellular ice crystals which can rupture cell membranes (12). The thawing process is typically quick, as microbial death is commonly associated with slow thaw rates. Though cryopreservation is a standard method for maintaining cellular organisms, it has rarely been utilized for the preservation of algal viruses.

One eukaryotic algal virus cryopreservation protocol is in existence. It was developed for HaV, a dsDNA virus that infects the red tide forming dinoflagellate *Heterosigma akashiwo* (13). Researchers investigated a combination of cryoprotectants and storage temperatures with the highest recovery (8.3% of infectious virus) employing flash freezing of HaV particles suspended in 20% DMSO. This protocol has been adapted for a handful of other algal viruses with viable recovery ranging from < 1% to 27% (14–16). The typical low recovery in these procedures is likely due to physiological differences between viruses and cells including differences in permeability, osmolarity tolerance, and toxicity to osmoprotectants. It is also clear that these protocols deviate from the standard method which controls the cooling rate; to our knowledge this has not been tested as a matter of improving virus particle survival. Owing to these complications, we decided to take a new approach by investigating cryopreservation recovery and stability of actively infected, cell-associated algal viruses.

Chloroviruses are large (> 300 kb), dsDNA viruses in the family *Phycodnaviridae* (17). They are members of the proposed order the Megavirales (18), also known as “giant” viruses, and remain the best characterized algal-virus system to date. Isolated in the early 1980’s (7), the prototype chlorovirus *Paramecium bursaria Chlorella virus 1* (PBCV-1) has been maintained through serial propagation on its host, *Chlorella variabilis* NC64A. PBCV-1 is inactivated by freezing, though other closely related virus strains, including other chloroviruses, persist through freeze/thaw events (19, 20). As a great deal of research has centered on PBCV-1, including genomics (21), transcriptomics (22, 23), and proteomics (21), it is important to develop a successful cryopreservation protocol for this strain that may serve as a model for preserving algal viruses. There are several reports of cryopreservation techniques for eukaryotic algae (24–28) which might be adapted for the preservation of actively replicating chloroviruses.

Here, we tested the cryo-potential of chlorovirus PBCV-1 using a protocol that yielded consistent recovery (~50% viable cells) of four strains of algae over 15 years: *Chlorella vulgaris* C-27, *Chlorella vulgaris* M-207A7, *Nannochloropsis oculate* ST-4, and *Tetraselmis tetrathlele* T-501 (29). Owing to the close relationship between *C. vulgaris* and *C. variabilis*, as well as the consistent results across unique algae, we elected to determine if these results could be recapitulated in PBCV-1. To test this, we attempted cryopreservation of both the virus particle as well as the virus replicating in its host.

## Materials and methods

### Virus particle cryopreservation

*Chlorella variabilis* NC64A was infected with PBCV-1 during mid-logarithmic growth at standard culturing conditions (25°C; continuous light exposure at 30μEin/m^2^/s) using Modified Bold’s Basal Medium (30). Following complete lysis, the viral lysate was pre-filtered through a sterile, 0.45 μm polycarbonate syringe filter and titered by plaque assay (31, 32) for initial infectivity assessments. Cryoprotectant choice was guided by Nakanishi *et al*. (29), in which a combination of 5% DMSO (v/v), 5% ethylene glycol (v/v), and 5% proline (w/v) was found to consistently produce the highest algal recoveries. Stock solutions of each cryoprotectant were made at a concentration of 30% with sterilized Milli-Q water and combined in a 1:1:1 ratio to yield a final concentration of 10% for each compound. For virus particle cryopreservation, 1 mL of PBCV-1 lysate (7.82x 10^8^ plaque forming units (PFUs) per ml) was added to 1 mL of ice-chilled cryoprotectant solution contained in a 2-mL cryovial. The cryovials were incubated on ice for 45 min, then transferred to a freeze-rate controlled container (Mr. Frosty, Thermo Fisher Scientific Inc., USA) filled with isopropanol for overnight incubation at −80° C. The next morning, cryovials were transferred to a −150° C freezer. At the designated recovery times, vials were removed from the freezer and set in a 40° C water bath. After thawing, the samples were serially diluted ten-fold in 50 mM Tris-HCl (pH = 7.8) and virus infectivity was determined by plaque assay (31). Virus viability was calculated as a percentage by comparison to the initial lysate titer before cryopreservation. Long-term experiments assessed the stability of virus infectivity in lysate stored at −150° C.

### Infected Chlorella cryopreservation

Chlorovirus PBCV-1 was propagated as described above and titered to obtain infectious PFUs/ml. This virus lysate was used to infect late-logarithmically growing *C. variabilis* NC64A at an M.O.I. of 5, at which point infected cultures were returned to standard incubation conditions. At 1, 10, 30, 60, 120, 180, 240, 300, and 360 min post-infection (PI), 1 mL aliquots of infected cells was mixed with 1 mL of ice-chilled cryoprotectants [final concentration: 5% DMSO (v/v), 5% ethylene glycol (v/v), and 5% proline (w/v)] in duplicates. The mixture was incubated on ice for 45 min, then transferred to a freeze-rate controlled container (Mr. Frosty, Thermo Fisher Scientific Inc., USA has a −1C/min cooling rate) filled with isopropanol for overnight incubation at −80° C. The next morning, cryovials were immediately transferred to a −150° C freezer. At the designated recovery times, vials were removed from the freezer and placed in a 40° C water bath. After thawing, the infected cells were pelleted in a Sorvall Legend RT Benchtop Centrifuge at 3,700 rpm (~3,000 rcf) for 10 min: (free virus requires higher speeds for pelleting). Cell pellets were re-suspended in 2 mL of 0.01M HEPES solution (pH = 6.5). Suspensions were immediately diluted and plaque assayed, plating late-infection treatments first. Viability was determined as a percentage of the pre-frozen cellular concentration (3.57 x 10^6^ cells/mL), as only surviving infected cells would be capable of producing plaques. Long-term experiments were conducted in the same manner, though only time points 10, 180, and 240 min PI were collected and assayed. The complete step-by-step method can be found at protocols.io (33).

## Results

Following the cryopreservation procedures of other algal virus researchers (13–16), we investigated the cryo-potential of the PBCV-1 lysate. Cryoprotectant alone treatments elicited a lethal effect: ~87% of the infectious virus particles were inactivated in the presence of these chemicals following 24 hr exposure at 4° C. Given this effect, we decided to freeze PBCV-1 lysate at −150° C without any cryoprotectants. This resulted in ~2.5% recovery of the infectious virus population, which was stable for storage periods of up to one year (Fig 1). Seeing room for improvement, we tested the cryo-potential of PBCV-1 in an infected, cell-associated state.

**Fig 1.**
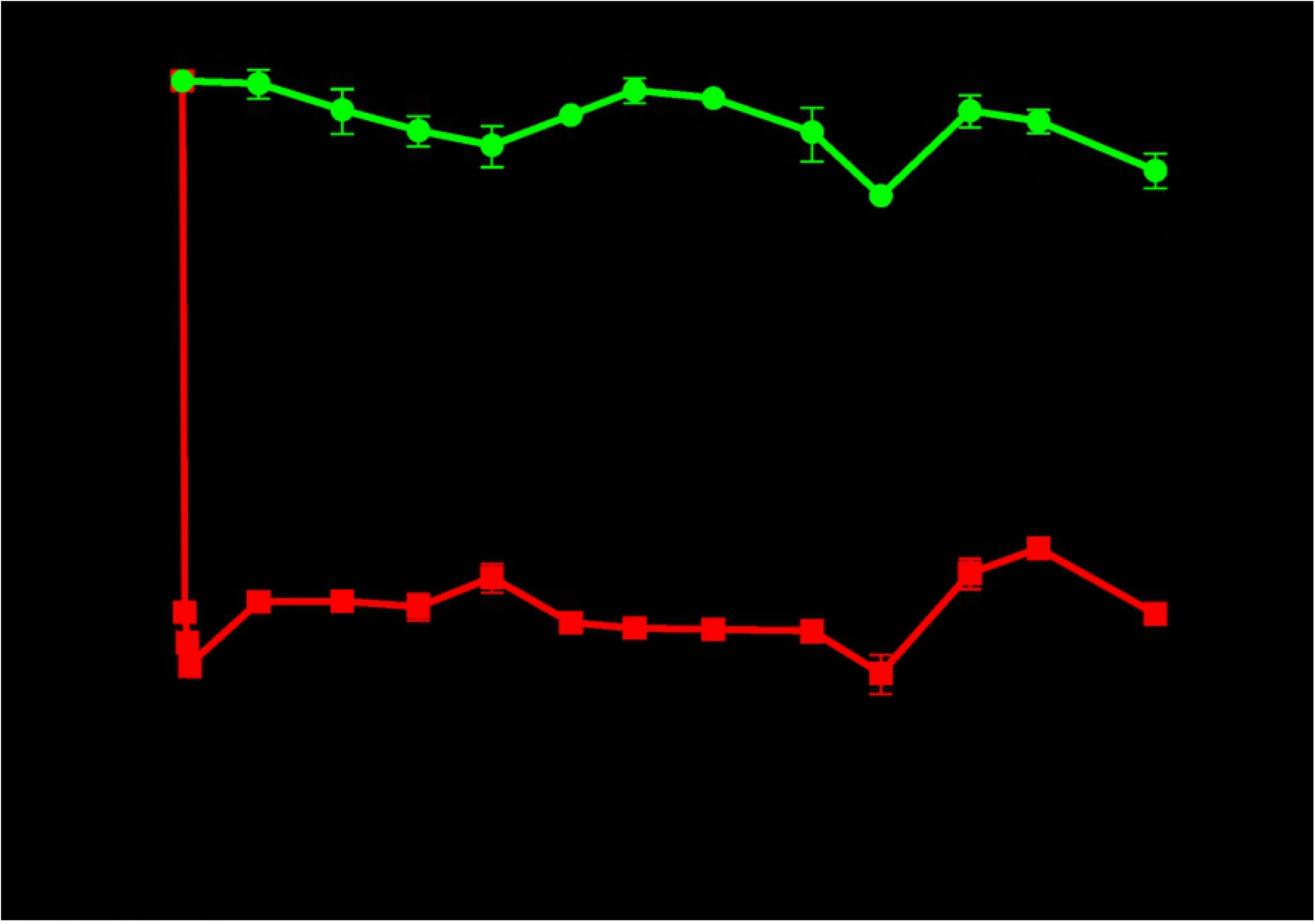
Cryo-stability of the PBCV-1 particle. Viability of chlorovirus PBCV-1 was determined by plaque assaying viruses that had been stored as lysates either at 4°C or −150°C. Green circles represent lysate stored at 4° C, while red squares denote lysate stored at −150° C. Error bars are represented as the standard deviation of biological and technical replicates.

The PBCV-1 replication cycle requires about 6-8 h to release nascent virus particles (34). Post-infection sampling times for cryopreservation (10, 30, 60, 120, 180, 240, 300, 360 min PI) followed similar sampling strategies used in PBCV-1 transcription studies (22, 23). Specifically, these time points were collected across distinct physiological phases in the PBCV-1 lifecycle and thus represent likely unique conditions for cryopreservation. Following 24-h storage of cryopreserved, infected cells, we found that late stages of infection were more conducive to virus survival than early stages (Fig 2). Thus, we followed cryo-stability for one year in one early (10 min PI) and two late infection stages (180 and 240 min PI) (Fig 3). Small day-to-day fluctuations in virus titers were common, but were typically consistent among treatments, suggesting human error. Despite these fluctuations, the lysate control, 180-min, and 240-min PI treatment yielded an acceptable relative standard deviation (RSD) across all recovery assessments, indicating cryo-stability (Table 1). Cryo-stability was not observed in the 10 min PI samples (Table 1). In comparison to virus lysate cryopreservation, the cell-associated method yielded significant improvement in survivorship for the optimal 240-minute treatment (24.9 – 30.1 fold increases).

**Fig 2.**
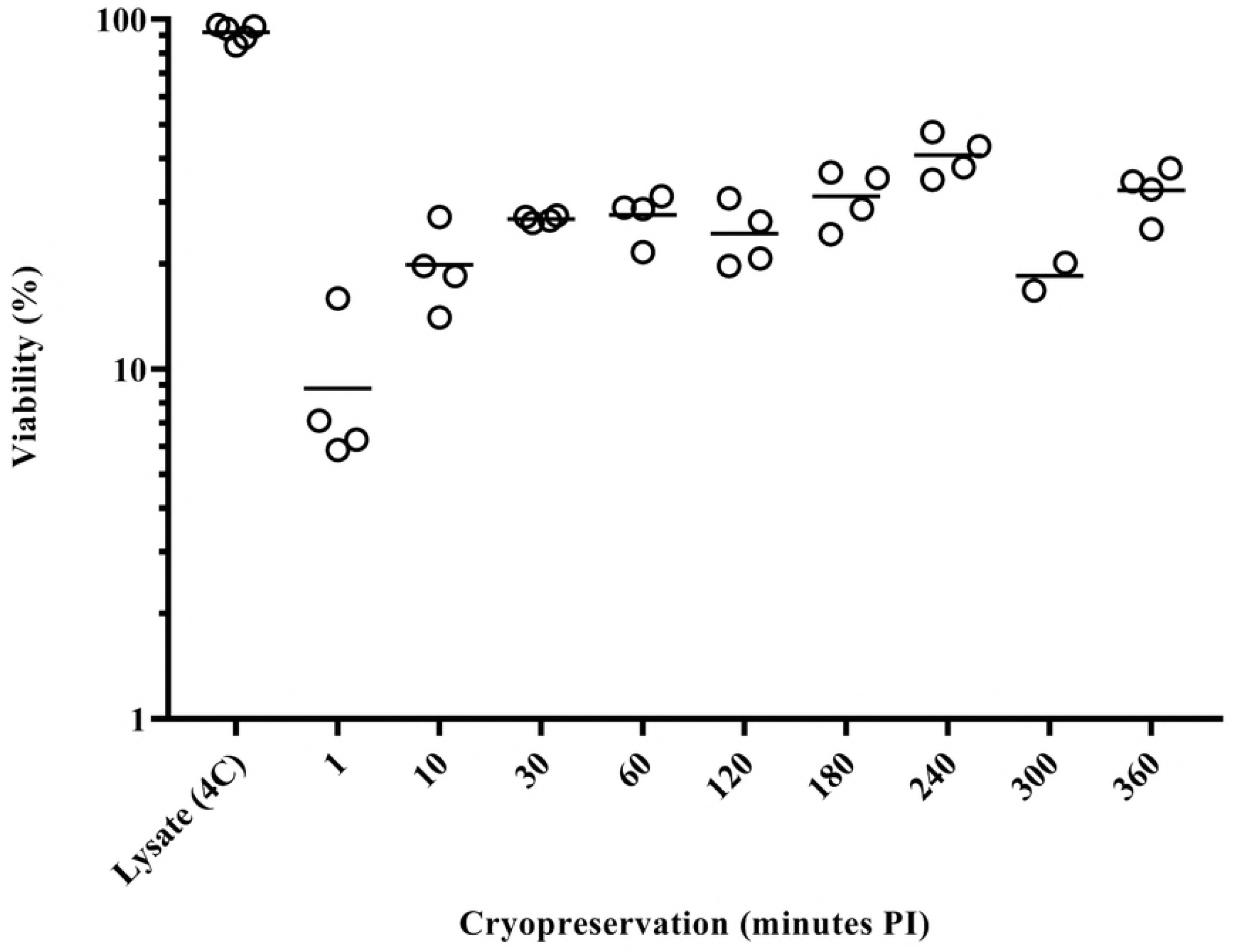
Recovery of infectious PBCV-1 frozen at various times after PBCV-1 infection of the *C. variabilis* NC64A host. Viability of chlorovirus PBCV-1 was assayed by monitoring plaque formation of cell-associated viruses that were collected at different times during an active infection cycle of the NC64A host. Open circles denote replicate plaque titers, with the average represented by the solid line.

**Fig 3.**
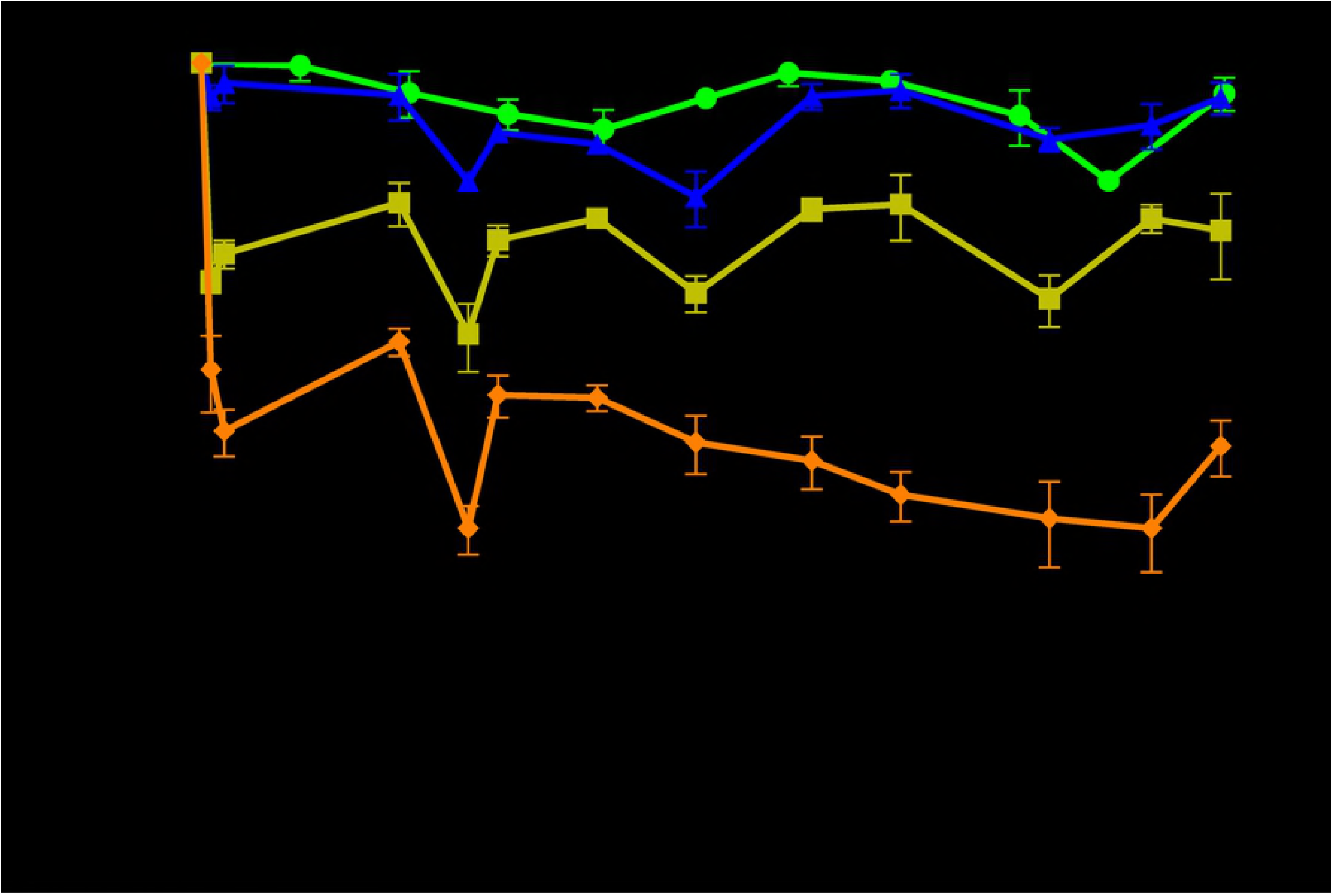
Long-term cryo-stability of PBCV-1 frozen in host cells at various times after infection of its host *C. variabilis* NC64A. Infectious chlorovirus PBCV-1 was monitored by plaque assay in virus lysate stored at 4° C (green circles) and in cryopreserved, PBCV-1-infected host cultures. Blue triangles, yellow squares, and orange diamonds represent virus viability following storage of infected cells cryopreserved after 240, 180, and 10 minutes PI. Error bars represent the standard deviation among biological and technical replicates.

**Table 1.** Statistical assessment of PBCV-1 infectivity across storage treatments for ~1 year.

| Treatment | N | Average | SD | RSD | 95%CI |
| --- | --- | --- | --- | --- | --- |
| Virus Particle Stock (4° C) | 67 | 75.1 | 16.9 | 22.5* | 71.1 - 79.2 |
| Virus Particle Stock (-150° C) | 124 | 2.53 | 0.61 | 24.0* | 2.42 – 2.64 |
| Cell-associated virus 10 minutes PI (-150° C, +CPA) | 79 | 7.56 | 3.38 | 44.7 | 6.81 – 8.31 |
| Cell-associated virus 180 minutes PI (-150° C, +CPA) | 82 | 31.9 | 10.9 | 34.2* | 29.5 – 34.3 |
| Cell-associated virus 240 minutes PI (-150° C, +CPA) | 82 | 69.5 | 16.5 | 23.8* | 65.9 – 73.0 |

+CPA, cryoprotectants present as described in materials and methods section. Asterisks denote an acceptable RSD (*i.e.*, Coefficient of Variation) for plaque assays based on a 35% threshold used in bacterial plating standards set from chapter 1223 by the U.S. Pharmacopeia and National Formulary (35).

## Discussion

The current maintenance strategy for chloroviruses involves serial propagation on the alga host followed by lysate storage at 4°C. Chloroviruses are stable under these conditions, but stocks can become contaminated by bacteria and rapidly degrade. Moreover, serial propagation of viruses, even if done only a few times a year, can promote genetic drift and result in deviation from wild-type status. This is concerning for all virus types, though RNA viruses, which have the fastest mutation rates, would be most susceptible (9, 36). Beyond considering spontaneous, replication-associated errors, chloroviruses encode putative enzymes involved in genomic rearrangements. For example, GIY-YIG mobile endonucleases and an IS607 transposon may be involved in insertions/deletions and/or gene loss/duplications observed in genomic comparisons of chloroviruses (37, 38). Thus, maintenance of wild-type strains is important for consistency between experiments. Virology labs could follow the microbial culture collection strategy, which typically uses a cryo-banking/seed-stock system for the dissemination of microbial specimens. The purpose of the seed-stock system is to minimize serial propagation of microbiota. The American Type Culture Collection (ATCC) suggests that consumers transfer their cultures no more than five-times after propagation from the thawed culture collection stock. Though a seemingly strict standard, it is not difficult to imagine the consequences of violating this. For example, the United States Pharmacopeia and National Formulary requires test organisms to be maintained this way for routine antibiotic efficacy screens, and non-compliance can undermine therapeutic treatment (35). Although there is no direct clinical link to maintaining algal viruses this way, the logic is consistent with any research requirements. The cryopreservation protocol described here can help researchers better set up these cryo-banking/seed stock systems.

Standard cryopreservation techniques are not designed for the unique structure and physiology of virus particles. Indeed, cryoprotectants are classified by their permeability across cell membranes, which often coincides with their molecular weight (24). Smaller compounds, such as ethylene glycol and DMSO, are considered penetrating cryoprotectants, while larger compounds (*e.g*. amino acids; L-proline) are typically non-penetrating. That said, the exclusion size threshold has not been established for most viruses so it is not clear which, if any of these compounds penetrate the viral capsid. It is generally thought that virus capsids are permeable to water and ions, though the latter diffuses much slower; this mechanism has been used to osmotically rupture capsids (39, 40), including PBCV-1 (41). The final cryoprotectant solution used for PBCV-1 lysate cryopreservation has an estimated osmolarity of ~150 mOsmoles/L, which is comparable to the storage buffer used for this virus. In light of this, we propose that the lethal effect the cryoprotectants have on the PBCV-1 particle is not the result of osmotic stress, and that inactivation instead occurred by toxicity of cryoprotectants or oxidative stress. This would be consistent with viruses not being metabolically active and therefore unable to repair damage caused by this treatment. It is also consistent with the observation that Mimivirus, a giant virus relative which also contains an internal lipid membrane, is said to be inactivated by lipophilic compounds such as DMSO (42). That said, DMSO is often used as a stabilizer for freezing of enveloped virus particles (43). This discrepancy may be due to unique properties between external and internal membranes, or even system differences between animal and plant viruses, which imparts resistance in some cases over others. Regardless, the mechanism of inactivation may be better ascertained by looking at survivorship of virion particles via epifluorescent microscopy, flow cytometry (44–46), or using bioassays to quantify oxidative stress.

Although the algal cell is in a sub-optimal physiological state during infection, it is apparently robust enough to survive and maintain an active infection during cryopreservation. That said, fewer infectious virus were recovered when the cell was cryopreserved during early infection stages. This might be explained by differences in adsorption rates and synchronicity of infection, resulting in fewer infected cells at the start of the experiment. Most, if not all cells are infected at the later stages of infection (3-4 hr PI). Regardless of any differences in synchronicity, the algal cell will be completely arrested during cryopreservation, and will only continue the infection cycle after thawing. Internal, mature viruses that have not yet lysed their host cell might still be inactivated by cryoprotectants, thus reducing viral burst size, but our experiments did not account for this.

The general classification of cryoprotectants based on membrane permeability is consistent in the infected cell treatment. Although the *C. variabilis* NC64A genome encodes a secondary active transporter for the uptake of proline, radio-labeled solute uptake experiments revealed that PBCV-1 infection abolishes its activity (48). With that in mind, the tonicity of the cryoprotectant mixture would equate to ~90 mOsmoles/L, as only DMSO and ethylene glycol are penetrating, and many of the components in the MBBM media would be spent by late-logarithmic growth. This concentration is comparable to buffers routinely used in our lab for handling *C. variabilis* (40 mOsmoles/L), so there is little concern of osmotic stress. The chances of osmotic stress were also low considering the consistent success associated with this cryopreservation formula across eukaryotic algae, including two *Chlorella* spp. (29). Our results are likely applicable to any algal virus whose host can be cryopreserved. That said, we expect that researchers may still have to adjust their cryoprotectant mixture to account for system differences related to osmolarity tolerance and cryoprotectant toxicity. There has also been research indicating that axenicity impacts cryopreservation survival in microalgae. In this light, it is possible that the bacterial community produces secondary metabolites which promote survival (49). In another scenario, organisms with psychrophilic tendencies might be adapted to freeze situations and cryoprotectant additives may not be necessary.

The goal of this study was to develop a long-term cryopreservation method for chlorovirus PBCV-1, but there are also interesting ecological implications of this research. Recent metagenomic and isolation efforts indicate that giant viruses of microeukaryotes (e.g., *Phycodnaviridae* and *Mimiviridae*) are widely distributed in nature (50, 51), but it is not well understood how these viruses persist in the environment. Freezing events represent a potential mechanism of inactivation for some algal viruses, though chlorovirus ATCV-1 is stable during these conditions (19). In two other studies, a closely related giant virus of the family *Mimiviridae* (52), as well as a second giant virus in the family *Molliviridae* (53), were revived from 30,000 year old permafrost. Both of these viruses were revived using *Acanthamoeba spp*., one of the main hosts for many giant viruses. That said, there have been questions about whether *Acanathamoeba* and other protists used for laboratory viral propagation are the natural or primary hosts of these ancient viruses (54). Although these viruses might be able to withstand freezing temperatures on their own, the results of this study suggest that a natural host might serve as a better vehicle for surviving freezing. Indeed, many microbes produce natural cryoprotectants (*e.g*. L-proline, trehalose, betatine, etc.) or encode machinery to transport these osmoprotectants into the cell. Following this thought process, it is possible that environments containing frozen, infected cells might contain naturally cryopreserved algal-virus systems. These systems may be deciphered following advances in single-cell sorting and sequencing techniques. Indeed, a similar approach has been successfully utilized to identify and sequence single virus genomes in the ocean (55). Though this latter study sorted virus particles, flow-cytometry sorting of viral infected cells may be achieved using fluorescent probes specific for viral marker genes (*e.g*., major capsid protein) or dyes to detect viral-induced host phenotypes (*e.g*., membrane blebbing). As a proof of concept, viral genetic sequences from Siberian permafrost could be used to probe for still frozen viral-infected host cells, thereby testing the natural host range of these viruses.

To our knowledge, this is the first report of successful cryopreservation of a eukaryotic algal virus during its infection cycle. We expect that respective cellular hosts will provide more suitable physiological conditions for cryopreservation and storage of algal viruses that infect eukaryotic algae. We also recommend that laboratories working with algal viruses establish cryopreserved seed-stock systems to better preserve wild-type controls for future experimentation, especially in lieu of future modification of these viral systems.

## Acknowledgements

We thank Professor David A. Hutchins (University of Southern California) for his thoughtful discussion on this manner over 2 decades ago. This work was support by grants from the National Science Foundation (NSF-OCE 1829641) and the Gordon & Betty Moore Foundation (#4971) to SWW.

